# Conjugation-based genome engineering in *Deinococcus radiodurans*

**DOI:** 10.1101/2021.10.13.464295

**Authors:** Stephanie L. Brumwell, Katherine D. Van Belois, Daniel J. Giguere, David R. Edgell, Bogumil J. Karas

**Author notes:** **Corresponding Author** Bogumil J. Karas.

## Abstract

*D. radiodurans* has become an attractive microbial platform for the study of extremophile biology and industrial bioproduction. To improve the genomic manipulation and tractability of this species, the development of tools for whole genome engineering and design is necessary. Here, we report the development of a simple and robust conjugation-based transformation system from *E. coli* to *D. radiodurans* allowing for the introduction of stable, replicating plasmids expressing antibiotic resistance markers. Using this method with nonreplicating plasmids, we developed a protocol for creating sequential gene deletions in *D. radiodurans* by targeting re-striction-modification system genes. Importantly, we demonstrated a conjugation-based method for cloning the large (178 kb), high G+C content MP1 megaplasmid from *D. radiodurans* in *E. coli*. The conjugation-based tools described here will facilitate the development of *D. radiodurans* strains with synthetic genomes for biological studies and industrial applications.

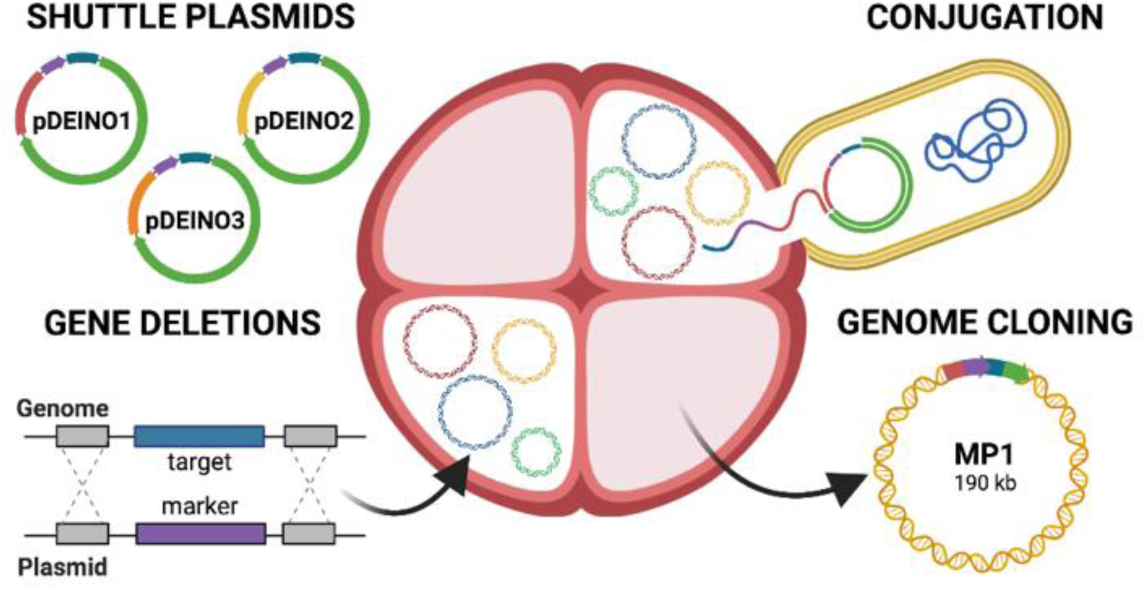

## INTRODUCTION

*Deinococcus radiodurans* is a polyextremophile bacterium that is well-known for its resistance to damage caused by ionizing and UV radiation^1,2^, desiccation^3^, and the vacuum of space^4–6^. Its survivability has been the focus of many studies and has primarily been attributed to efficient protection against proteome degradation, ensuring cellular recovery and repair of damaged DNA^7^. These characteristics make *D. radiodurans* an attractive chassis for biotechnology as it can withstand the physiochemical stress of industrial processes that traditional model organisms, such as *Escherichia coli* and yeast, would be unable to survive. Currently, this organism is used for biotechnological applications including bioremediation^8,9^ and the production of small molecules^10^ and unique pigments^11,12^, but it has even greater potential applications that could be elucidated by genome engineering.

*D. radiodurans* R1 was first sequenced using whole genome shotgun sequencing in 1999 by White *et al*., which revealed that its genome was composed of two chromosomes (2.6 Mb and 412 kb), a megaplasmid (MP1, 177 kb) and small plasmid (CP1, 46 kb), containing 67% G+C content on average^13^. Recent PacBio sequencing of this strain resulted in a high quality genome that identified both large insertions and single nucleotide polymorphisms (SNPs) when mapped to the original sequence^14^. Engineering in this organism is progressing with the available sequence data and expanding genetic toolkit including shuttle plasmids^15–17^, selection markers^18–20^, characterized promoters for inducible and tunable gene expression^21,22^, as well as a strategy for generating gene deletions using a Cre-lox system^23^. In order to employ engineering strategies, it is crucial to have a method of delivering DNA to the host which can currently be achieved in *D. radiodurans* by natural transformation, chemical transformation and electroporation^24^. However, in most cases these methods require large amounts of DNA, likely as a result of active restriction-modification systems known to digest exogenous DNA. Furthermore, the polyploid nature of *D. radiodurans* can make genetic modification more difficult as there can be up to 10 copies of its genome^25,26^. Thus far, the transfer of large genomic regions or whole chromosomes has not yet been demonstrated in this species but would considerably improve the ability for large-scale engineering.

Although *D. radiodurans* was one of the first bacteria to be sequenced, tools to enable whole genome engineering have yet to developed due to the aforementioned drawbacks. These tools could provide new insights into the minimal set of genes required for life in extreme environments and allow for the production of improved industrial strains, using similar methods as the genomic recoding or streamlining demonstrated in *Mycoplasma mycoides*^27,46^ and *E. coli*^28,47^. Large-scale genetic manipulation typically involves the generation of synthetic DNA fragments or genomes in host organisms, like *E. coli* or yeast, followed by transfer to destination cells via transplantation^27^, bacterial conjugation^28^, or electroporation^29^. Towards the goal of whole genome engineering, methods for DNA delivery to *D. radiodurans* must be improved first.

Conjugation could potentially address this bottleneck as it has been shown as a simple and efficient method for direct DNA transfer between bacteria and from bacteria to eukaryotes. For industrial applications in the field of synthetic biology, conjugative systems often utilize a two-plasmid approach. The conjugative plasmid contains all genes needed to form a pilus between the donor and recipient organisms, as well as those needed to mobilize the cargo plasmid. The cargo plasmid can contain any genetic construct of interest, accompanied by an origin of transfer (*oriT*) essential for proteins encoded on the conjugative plasmid to initiate and facilitate DNA transfer to a recipient cell. Conjugation can also streamline the genetic manipulation of organisms as it eliminates the need for high yield DNA extraction and has proven to be a versatile method for moving large plasmids to both bacterial and eukaryotic organisms^28,30–35^.

Here, as the first step towards genome-scale engineering in *D. radiodurans*, we have developed a conjugation protocol to transfer replicating and nonreplicating plasmids from *E. coli* directly to *D. radiodurans* and determined the conjugation frequency. We demonstrated a conjugation-based method for creating targeted gene deletions, replacing four restriction-modification system genes with antibiotic resistance markers. Finally, we cloned the whole 178 kb MP1 megaplasmid from the *D. radiodurans* genome in *E. coli* using conjugation-based plasmid integration.

## RESULTS AND DISCUSSION

We developed a conjugation protocol as a direct method of DNA transfer from *E. coli* to *D. radiodurans* (Figure 1). To achieve this, we first constructed a replicating plasmid, pDEINO1, by assembling a codon-optimized chloramphenicol acetyltransferase gene (*cat*) and an origin of replication^17^ for *D. radiodurans* into the pAGE3.0^30^ multi-host shuttle (MHS) plasmid (Figure 2A). We used an *E. coli ΔdapA* donor strain^30^ to allow easy counter selection of the donor bacteria and wild-type *D. radiodurans* as the recipient. The donor strain harbors the conjugative plasmid pTA-Mob^36^, an RP4/RK2 derivative, and the ∼22 kb mobilizable plasmid, pDEINO1. Conjugation frequency was determined by the proportion of *D. radiodurans* colonies that formed on media supplemented with chloramphenicol to the number of colonies on nonselective media. The mean conjugation frequency from *E. coli* to *D. radiodurans* was 1.12 × 10^−5^ transconjugants per recipient (Figure 2B). When *E. coli* harboring pDEINO1, but lacking pTA-Mob, was used as a donor for conjugation there were no transconjugant colonies on selective plates. There were also no transconjugant colonies formed when the recipient was plated with no *E. coli* donor.

**Figure 1.**
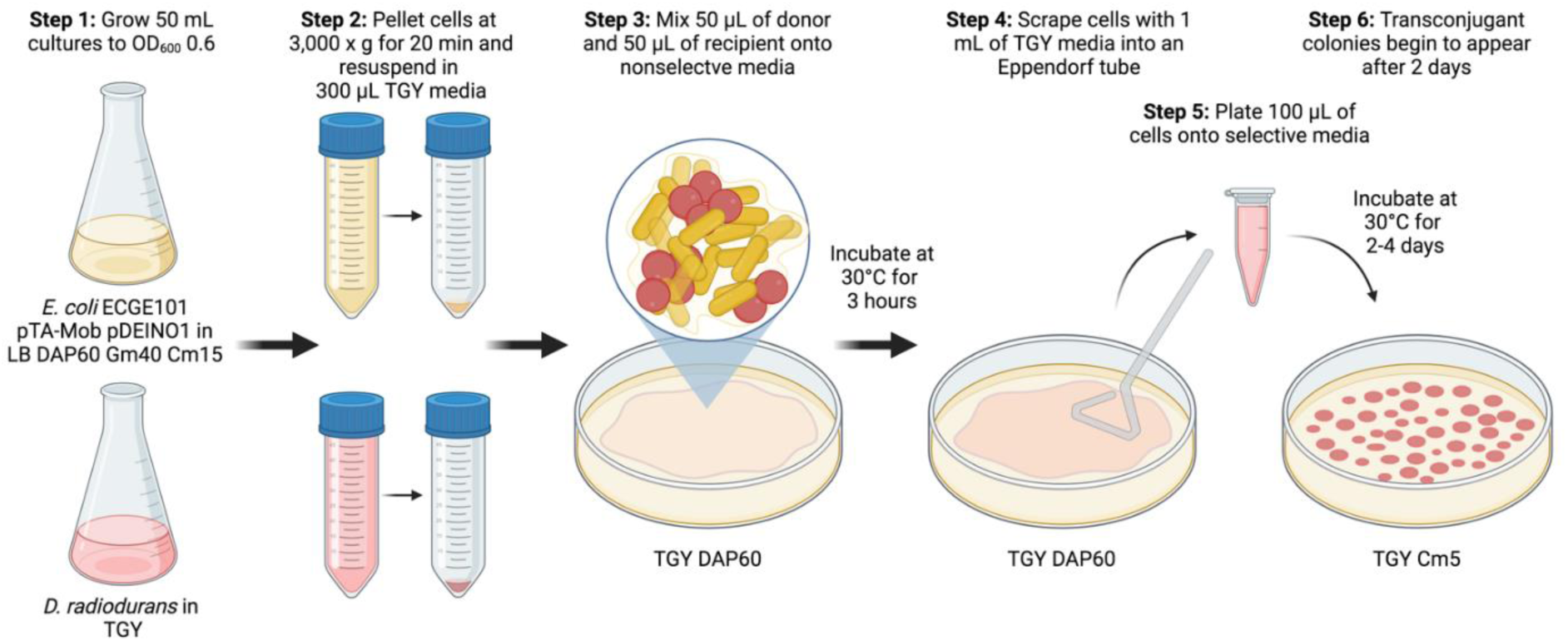
Flow chart of the *E. coli* to *D. radiodurans* conjugation protocol. Antibiotic concentrations are reported in μg mL^-1^. Created with BioRender.com.

**Figure 2.**
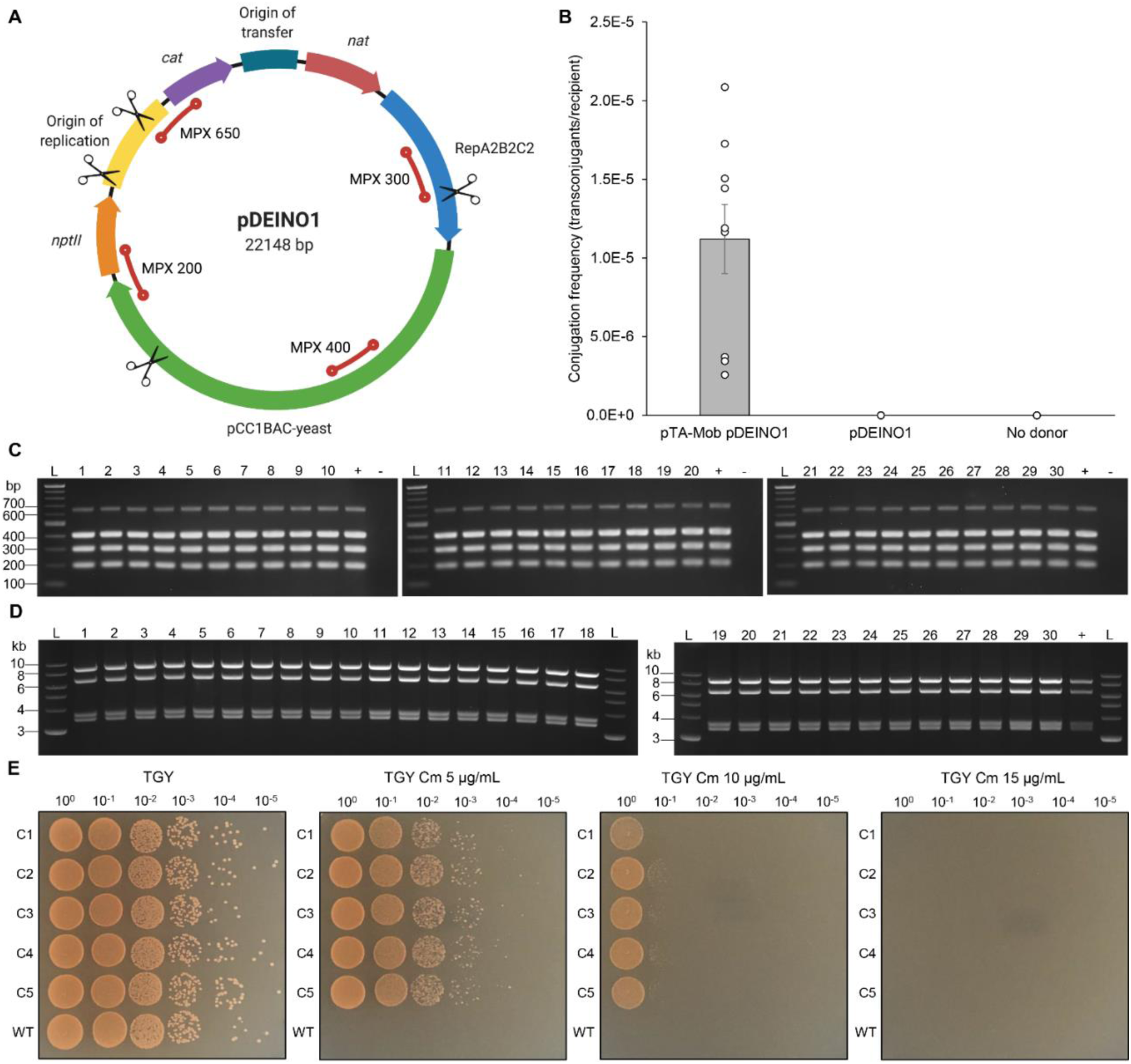
Demonstration of pDEINO1 plasmid conjugation from *E. coli* to *D. radiodurans*. **(A)** Schematic of the multi-host shuttle plasmid pDEINO1. *cat*, codon-optimized chloramphenicol resistance gene for *D. radiodurans. nat*, N-acetyltransferase. *nptII*, neomycin phosphotransferase II. RepA2B2C2, *S. meliloti* origin of replication. pCC1BAC-yeast, backbone for replication in yeast and *E. coli*. Red lines indicate the location and size (bp) of expected multiplex amplicons and EcoRI-HF cut sites for the diagnostic digest are represented as scissors. Created with BioRender.com. **(B)** Conjugation frequency of pDEINO1 from *E. coli* to *D. radiodurans* with *E. coli* donors harboring pTA-Mob and pDEINO1, only pDEINO1, or experiments performed without *E. coli* donors, only recipient cells. Data are shown as a bar graph with points representing individual technical replicates. Error bars are standard deviation of the mean. **(C-D)** Multiplex PCR and diagnostic restriction digest of 30 pDEINO1 plasmids from *D. radiodurans* transconjugants, following transformation and plasmid induction in *E. coli*. L, 2-log ladder, +, original pDEINO1 plasmid prior to conjugation. -, water. **(C)** Expected multiplex amplicon sizes are 200, 300, 400, and 650 bp. **(D)** Expected EcoRI-HF digest band sizes are 3149, 3450, 6548, and 8474 bp. **(E)** Phenotypic screening for chloramphenicol sensitivity of *D. radiodurans* transconjugants harboring pDEINO1 relative to wildtype *D. radiodurans* by serial dilution spot plates on TGY media and TGY media supplemented with increasing concentration of chloramphenicol. C1-C5, *D. radiodurans* transconjugant colonies. WT, wildtype *D. radiodurans*.

To confirm that the *D. radiodurans* transconjugants harbored pDEINO1, we isolated plasmids from 30 individual *D. radiodurans* transconjugant colonies. The DNA was transformed into *E. coli* Epi300, colonies were selected on media supplemented with chloramphenicol, and plasmids from 30 single colonies were induced to high copy number with arabinose. The plasmids were analyzed by multiplex PCR (Figure 2C) and diagnostic restriction digest (Figure 2D), both of which showed the expected banding patterns following gel electrophoresis. Additionally, sequencing analysis of plasmids from two transconjugant colonies showed no mutations when compared to the original pDEINO1 sequence. These results demonstrate that replicating plasmids can be successfully conjugated from *E. coli* to *D. radiodurans* with no gross rearrangements or point mutations. Furthermore, the codon-optimized chloramphenicol acetyltransferase gene is an effective plasmid-based marker for selection of *D. radiodurans* trans-conjugants. This strategy of using conjugation to deliver replicating plasmids circumvents the difficulties associated with transforming DNA into strains containing active restriction systems.

To determine the sensitivity of *D. radiodurans* to chloramphenicol, five transconjugant colonies were spot plated alongside the wild-type strain on varying concentrations of chloramphenicol (Figure 2E). We found that chloramphenicol at a concentration of 5 μg mL^-1^ inhibited the growth of wild-type *D. radiodurans*, while having only a small negative effect on transconjugants; however, any additional increase in chloramphenicol concentration significantly impacted transconjugant growth. Additionally, we demonstrated that the *E. coli cat* gene present on pDEINO1 was not able to provide resistance for *D. radiodurans* when introduced on pDEINO6 without the *D. radiodurans cat* gene (Supplementary Figure 1).

Plasmid loss is beneficial when employing an engineering strategy such as those dependent on the recycling of plasmids or that require plasmids temporarily for expression of an endonuclease or Cas9, for instance. To demonstrate that the strains could be cured of the plasmid, we performed a plasmid-loss assay on one transconjugant colony grown without selection in liquid media for approximately 40 generations. At 13-generation intervals, the culture was plated on media with and without chloramphenicol and 100 colonies from each treatment were restruck onto selective plates. We showed that when growing without selection, on average 44% of cells lost the pDEINO1 plasmids every 13 generations (Supplementary Table 1). Although horizontal gene transfer to *D. radiodurans* has been predicted from comparative genomics studies^37^, suggesting that conjugation to this organism has occurred naturally before, our results are the first demonstration harnessing conjugative machinery for delivery of replicating and nonreplicating plasmids in a laboratory setting.

In order to design plasmid-based systems utilizing this new conjugative delivery method and to increase its potential for engineering *D. radiodurans*, we tested four additional antibiotic markers. Previous studies have demonstrated the use of *nptII* (Kan^R^/Nm^R^), *aadA* (Spec^R^), *cat* (Cm^R^) and *aac(3)* (Gen^R^) to select for genomic integration of transgenes^38^. Therefore, we cloned *nptII, tetR/A, aadA1*, or *aacC1* (conferring resistance to kanamycin/neomycin, tetracycline, streptomycin, and gentamicin, respectively) down-stream of the codon-optimized chloramphenicol marker onto separate replicating plasmids (Figure 3A). These plasmids, called pDEINO1, pDEINO3, pDEINO4 and pDEINO5, were conjugated into *D. radiodurans* from *E. coli* and transconjugant colonies were selected on plates supplemented with chloramphenicol. Individual colonies harboring each of the different plasmids were grown in liquid media alongside wild type *D. radiodurans*, and 10-fold serial dilutions were spotted onto plates supplemented with the aforementioned antibiotics.

**Figure 3.**
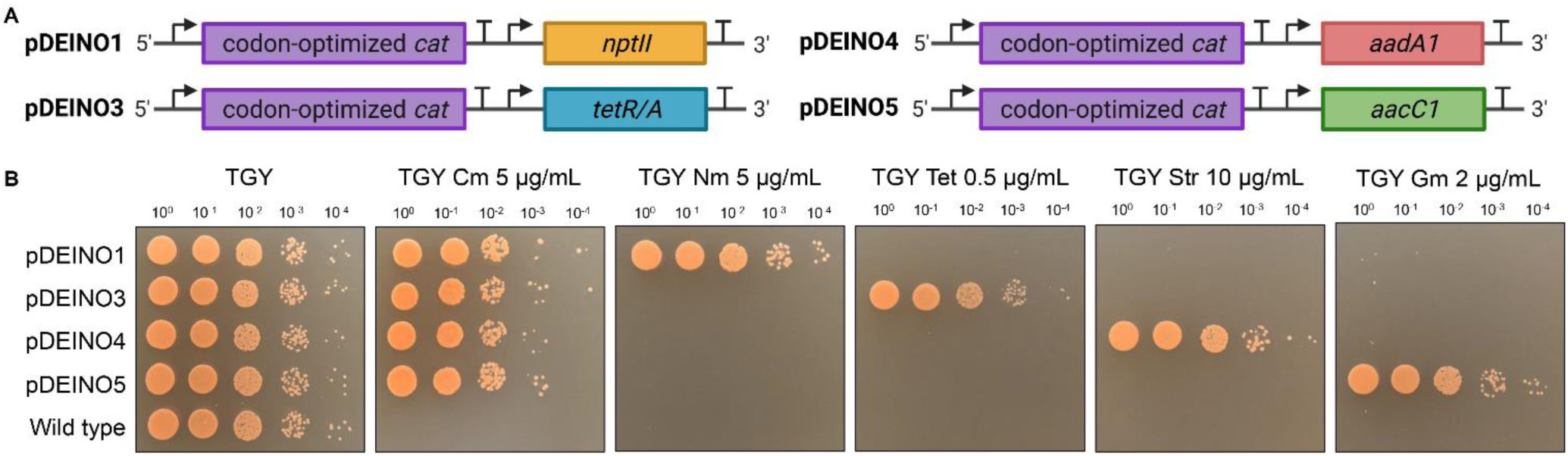
Test of selectable markers on replicating plasmids in *D. radiodurans*. **(A)** Diagram of the selective marker cassettes on pDEINO1, pDEINO3, pDEINO4 and pDEINO5 plasmids containing a codon-optimized *cat* gene for *D. radiodurans* coupled with *nptII, tetR/A, aadA1* or *aacC1*, respectively. Created with BioRender.com. **(B)** Serial dilutions of *D. radiodurans* harboring pDEINO1, pDEINO3, pDEINO4 and pDEINO5 alongside wild type *D. radiodurans* spot plated on TGY media and TGY media supplemented with chloramphenicol, neomycin, tetracycline, streptomycin and gentamicin.

As shown in Figure 3B, all strains harboring shuttle vectors conferred effective resistance to chloramphenicol and their respective antibiotic, while exhibiting sensitivity to the other three antibiotic supplements. Since *nptII* confers resistance to two antibiotics, the strain carrying pDEINO1 was also tested on media supplemented with kanamycin, though neomycin caused slightly less growth inhibition to transconjugants (Supplementary Figure 2). We observed that supplementing gentamicin at a concentration as little as 2 μg mL^-1^ sufficiently inhibited wild type *D. radiodurans* for our applications, although there was some breakthrough at the lowest dilution on the spot plates. Furthermore, changing the concentration by only 1 μg mL^-1^ significantly impacted the growth of *D. radiodurans* harboring pDEINO5. Increasing gentamicin concentration to 3 μg mL^-1^ severely inhibited growth, while decreasing the concentration to 1 μg mL^-1^ did not inhibit wild type *D. radiodurans* when compared to strains harboring the plasmid (data not shown). Our replicating plasmids were stably maintained in *D. radiodurans* with antibiotic selection. However, due to the sensitivity of *D. radiodurans* to antibiotics, tolerating concentrations as little as 0.5 μg mL^-1^ in some cases, the development of auxotrophic strains and compatible complementation plasmids could be beneficial for future research.

Gene deletions have previously been demonstrated in *D. radiodurans* using linear DNA fragments or plasmids containing regions homologous to the *D. radiodurans* genome flanking a selection marker^18^. However, integrated sequences were often heterozygous and could be deleted by intrachromosomal recombination events^26^. Targeted integration to create mutants became possible after the genome was sequenced^39^. This strategy has been used to introduce biosynthetic pathways into *D. radiodurans* for pollutant degradation or detoxification of mercury and toluene^8,9^. More recently, multiple knockouts of genes replaced by selection markers have been demonstrated by recombination of nonreplicating plasmid in *D. radiodurans*^38^, as well as a Cre-lox recombination system^23^.

With an efficient plasmid delivery method and a collection of selection markers to work with, we developed a conjugative plasmid-based protocol for generating gene deletions in *D. radiodurans*. We created a nonreplicating plasmid to include a construct of interest, in this case *nptII* or *tetR/A* resistance genes, between two regions of homology from the *D. radiodurans* genome flanking a gene targeted for deletion. We targeted four restriction-modification system genes in the *D. radiodurans* genome, some of which have been deleted in previous studies^40–42^. These genes include *Mrr* and ORF2230 located on chromosome 1, ORF14075 on chromosome 2, and ORF15360 on the MP1 megaplasmid (Figure 4A).

**Figure 4.**
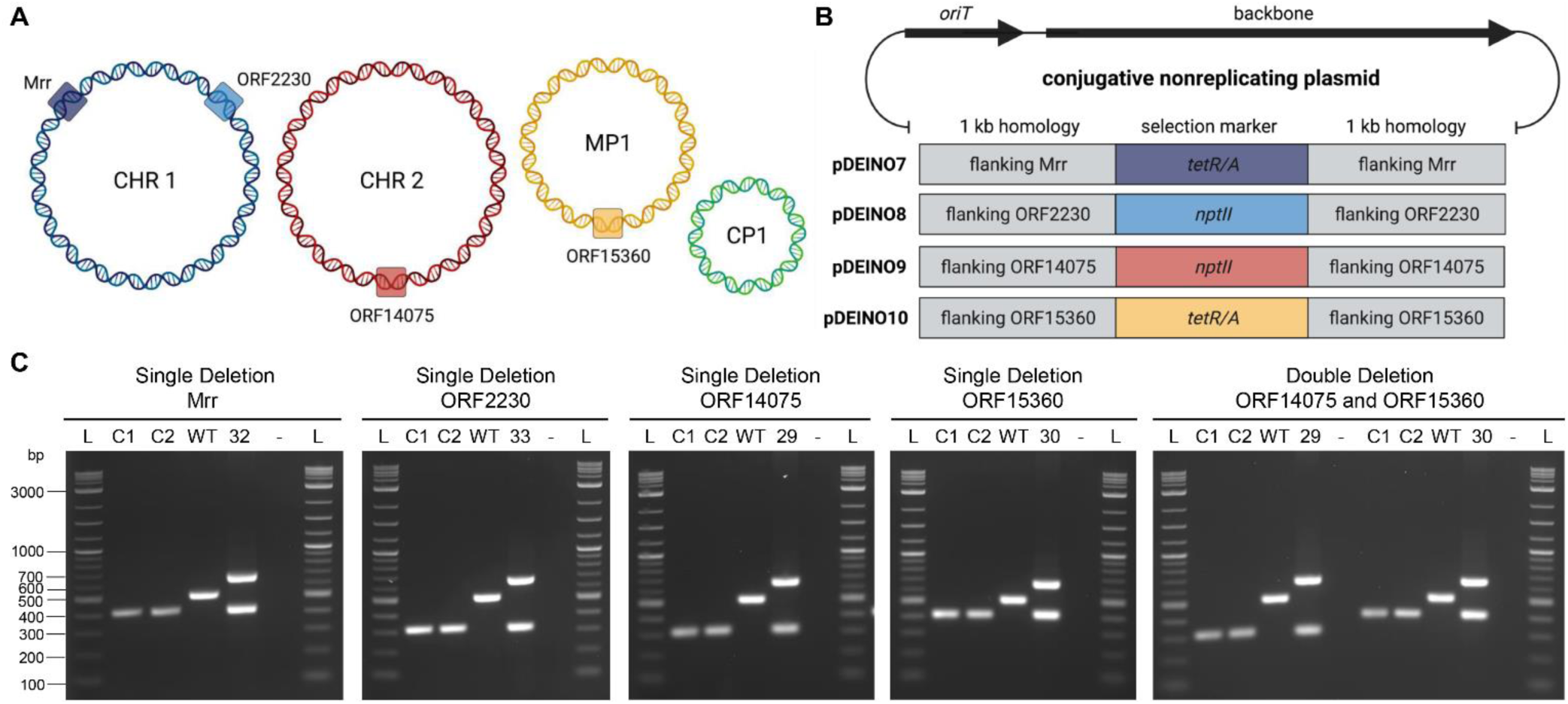
Demonstration of gene deletions in the *D. radiodurans* genome using recombination of conjugative suicide plasmids. **(A)** Diagram of *D. radiodurans* genome with the four restriction endonuclease targets indicated as coloured boxes. **(B)** Conjugative suicide plasmid map illustrating the plasmid components of pDEINO7, pDEINO8, pDEINO9 and pDEINO10, used to generate the *D. radiodurans* restriction endonuclease deletion strains. **(C)** Multiplex PCR analysis of two *D. radiodurans* transconjugant colonies for each restriction endonuclease knock out (C1, C2). Controls included wild type *D. radiodurans* genomic DNA(WT), the original conjugative suicide plasmid prior to conjugation (29, 30, 32, 33), and water (-). If present, amplicon sizes are: neomycin 311 bp, tetracycline 409 bp, wild type restriction endonuclease genes ∼500 bp, and the suicide plasmid backbone 645 bp. L, 2-log ladder. (**A**-**B)** Created with BioRender.com.

Prior to plasmid construction we resequenced our *D. radiodurans* R1 strain to ensure accurate homology sequences were incorporated into the plasmid designs. Genomic DNA was isolated using agar plug DNA isolation and was sequenced using Oxford Nanopore MinION sequencing technologies. Interestingly, we found that our plasmids did not contain the large insertions as reported in the PacBio sequencing results^14^, and the plasmid sizes more closely resembled the original sequence published by White *et al*.^13^.

All four nonreplicating plasmids were constructed with a backbone for replication and selection in *E. coli* and *Saccharomyces cerevisiae*, an *oriT* for conjugative transfer, and a deletion cassette containing a selectable marker – neomycin or tetracycline – between 1 kb regions of homology flanking the gene of interest (Figure 4B). Following conjugative delivery, transconjugants were selected with the respective antibiotic, DNA was isolated from *D. radiodurans* and screened for genomic gene deletions using multiplex PCR. Three amplicons could be observed in the multiplex, which reside within the following elements: selectable marker, plasmid backbone, and restriction endonuclease gene. Two transconjugants were screened for each single deletion event and PCR showed amplification of only the selectable marker, indicating successful integration of the selectable marker, deletion of the targeted gene, and loss of the plasmid backbone (Figure 3C). To show that this system could also be used to generate multiple deletions in the same strain, we performed conjugation using an ORF15360 transconjugant as the recipient and a donor *E. coli* harboring pDEINO7 targeting ORF14075. Transconjugants were selected on TGY media supplemented with both neomycin and tetracycline and their DNA was screen using the same multiplex PCRs as were used for the single deletion, confirming that both ORF15360 and ORF14075 genes were replaced with the tetracycline and neomycin markers, respectively, and the plasmid backbone was lost (Figure 4C). Successful deletion of multiple genes demonstrated that this technique could be implemented to sequentially delete genes involved in recognizing and digesting foreign or synthetic DNA in the cell in order to develop a restriction-free *D. radiodurans* strain.

To engineer *D. radiodurans* as an industrial platform organism, streamlined strains, such as those developed for *Mycoplasma mycoides* JCVI-syn3.0, may be developed by building synthetic genomes. Creating a minimal *D. radiodurans* cell will likely require the use of intermediate host organisms, such as *E. coli*, yeast, or *Sinorhizobium meliloti* to house the DNA and carry out efficient genetic modification and cloning^30,43,44^. To this end, we aimed to demonstrate that large DNA fragments from the *D. radiodurans* genome could be cloned in *E. coli*.

We built another nonreplicating plasmid called pDEINO2 and used our conjugation-based genome integration strategy to clone the native MP1 megaplasmid, 178 kb in length with a notable G+C content of ∼62%, from *D. radiodurans*. Mega-plasmids are large, extrachromosomal replicons that are commonly acquired through horizontal gene transfer. These replicons often provide beneficial, niche-specific traits to the host, such as antimicrobial resistance, symbiosis or metabolic pathways^45^.

To facilitate recombination and integration of the entire plasmid, pDEINO2 contained a1 kb region of homology to the middle of the *McrC* gene on the MP1 megaplasmid (Figure 5A). Following assembly, pDEINO2 was transformed into the Δ*dapA E. coli* donor strain and delivered to *D. radiodurans* via conjugation (Figure 5B). We confirmed recombination of pDEINO2 at the *McrC* gene by extracting DNA from transconjugants selected on TGY plates supplemented with chloramphenicol and performing a multiplex PCR amplifying within the non-replicative plasmid (Figure 5C, Supplementary Figure 3). Plasmid DNA was then transformed into *E. coli*, individual colonies were selected, induced to high copy number and isolated DNA was screened to verify the MP1 mega-plasmid had been transferred and propagated without compromising genomic integrity. The pDEINO2 plasmid was confirmed to be integrated at the correct location using two diagnostic digests (Figure 5D) and multiplex PCR that amplify regions of pDEINO2 and the *D. radiodurans* MP1 megaplasmid (Figure 5E). In particular, the presence of the 7733 bp band in the MfeI-HF digest was a strong indicator of plasmid integration at the *McrC* locus, as there is an MfeI site just outside the integration site and in the pDEINO2 backbone that creates a unique band size that would otherwise not be present in the wild type MP1 plasmid. The same is true for the 8112 bp band in the NheI-HF digest. Finally, we extracted total DNA from one *E. coli* clone harboring the pDEINO2-cloned MP1 plasmid and sequenced it using the Oxford Nanopore MinION.

**Figure 5.**
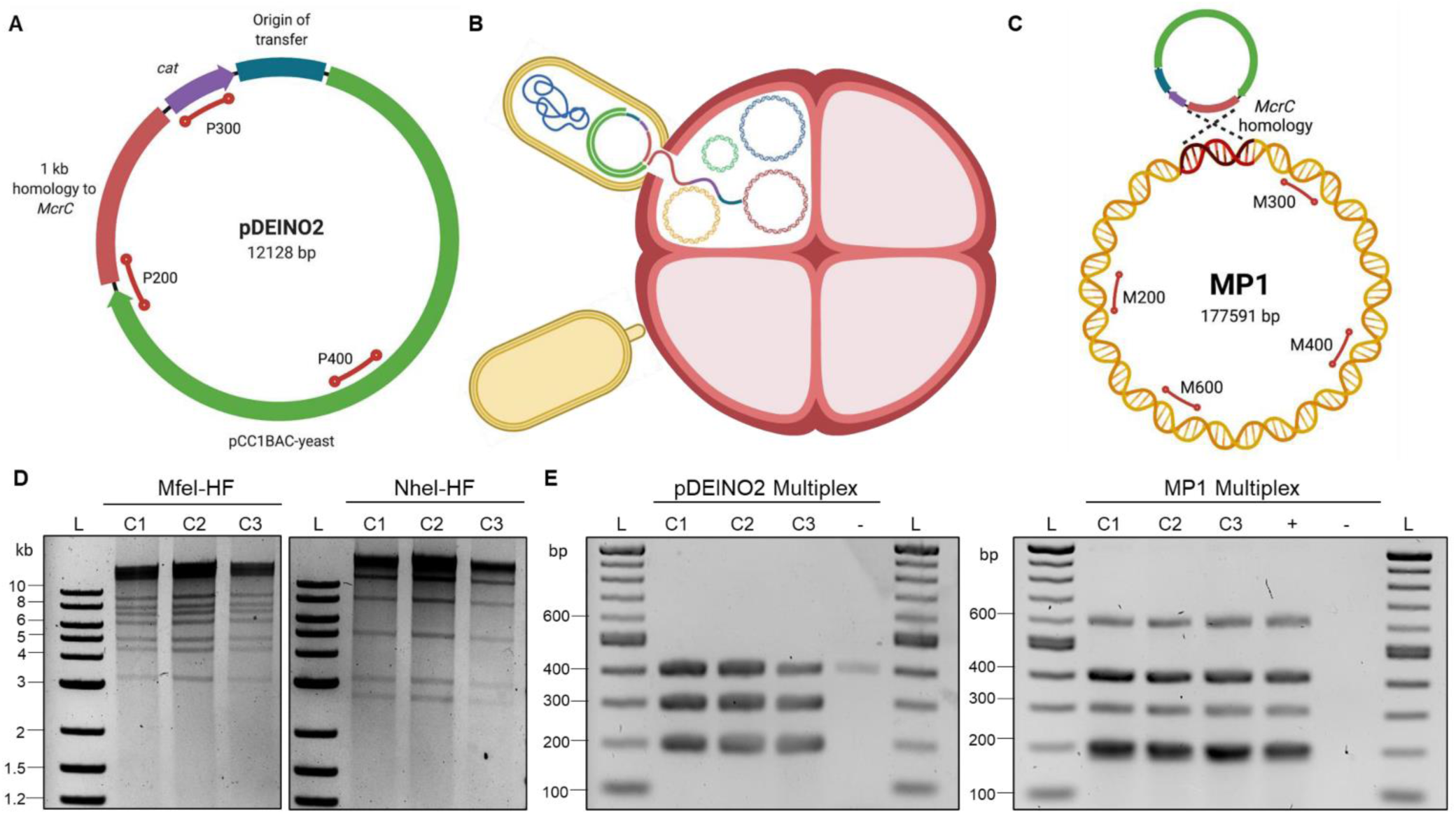
Cloning of *D. radiodurans* MP1 megaplasmid in *E. coli*. **(A)** Schematic of the pDEINO2 nonreplicating plasmid. Multiplex amplicon positions are indicated as red lines with the number indicating the amplicon size. **(B)** Diagram illustrating conjugation of the pDEINO2 plasmid from *E. coli* to *D. radiodurans*. **(C)** Diagram illustrating integration of pDEINO2 into the MP1 plasmid in *D. radiodurans* through recombination at the *McrC* gene. Multiplex amplicon positions are indicated as red lines with the number indicating the amplicon size. **(D)** Agarose gel of MfeI-HF and NheI-HF restriction digest of three cloned MP1 plasmids recovered from *E. coli* clones following induction. Expected visible band sizes (<10 kb) for the MfeI-HF digest are 325, 3186, 4338, 4929, 6170, 7049, 7733, and 8948 bp. Expected visible band sizes (<10 kb) for the NheI-HF digest are 2701, 3139, 5070, and 8117 bp. (**E)** Agarose gel of multiplex PCR of three cloned MP1 plasmids recovered from *E. coli* amplifying 200, 300 and 400 bp amplicons from the integrated pDEINO2 plasmid and 200, 300, 400 and 600 bp amplicons from the MP1 plasmid. L, 2-log ladder. +, *D. radiodurans* wild type genomic DNA. -, water. **(A-C)** Created with BioRender.com

The cloned MP1 plasmid was assembled and sequence data confirmed that pDEINO2 successfully integrated at the intended target site and the final plasmid size was ∼190 kb. Coverage for the cloned MP1 plasmid and sequencing depth for the genome indicated ∼7X multiplicity of the plasmid in *E. coli* following induction. These results indicated that large regions of the *D. radiodurans* genome, including entire replicons, can be effectively cloned and propagated in *E. coli*.

In summary, we have demonstrated conjugation from an *E. coli* donor as a method for DNA delivery to *D. radiodurans* and created several conjugation-based tools for genome engineering including gene deletions and whole chromosome cloning. Furthermore, we showed that *E. coli* can be a suitable host for building and maintaining large, high G+C content plasmids as demonstrated by cloning the MP1 megaplasmid from *D. radiodurans*. We anticipate that the genetic tools described here will rapidly advance the engineering process, particularly for modification of whole genomes, and allow for the development of *D. radiodurans* strains controlled by designer synthetic genomes for industrial, as well as foundational biology studies.

## METHODS

### Microbial Strains and Growth Conditions

*Deinococcus radiodurans* R1 (acquired from Dr. Murray Junop at Western University) was grown in TGY medium (5 g L^-1^ tryptone, 3 g L^-1^ yeast extract, 1 g L^-1^ potassium phosphate dibasic, and 2.5 mL 40% w/v glucose) supplemented with appropriate antibiotics (chloramphenicol 5 μg mL^-1^, neomycin 5 μg mL^-1^, tetracycline 0.5 μg mL^-1^) at 30°C. *Escherichia coli* (Epi300, Lucigen, Cat #: EC300110) was grown at 37°C in Luria Broth (LB) supplemented with appropriate antibiotics (chloramphenicol 15 μg mL^-1^). *Escherichia coli* ECGE101 (Δ*dapA*)^30^ was grown at 37°C in LB supplemented with diaminopimelic acid (DAP) 60 μg mL^-1^ and appropriate antibiotics (chloramphenicol 15 μg mL^-1^) and gentamicin 40 μg mL^-1^). *Saccharomyces cerevisiae* VL6-48 (ATCC MYA-3666: MATα his3-Δ200 trp1-Δ1 ura3-52 lys2 ade2-1 met14 cir^0^) was grown at 30°C in 2X YPAD rich medium (20 g L^-1^ yeast extract, 40 g L^-1^ peptone, 40 g L^-1^ glucose, and 200 mg L^-1^ adenine hemi-sulfate) supplemented with 80 mg L^-1^ adenine hemisulfate, or in complete minimal medium lacking histidine supplemented with 60 mg L^-1^ adenine sulfate (Teknova, Inc., Cat #: C7112) with 1 M sorbitol.

### Plasmid Design and Construction

All plasmids in this study (Supplementary Table 2) were constructed from PCR amplified DNA fragments assembled using a yeast spheroplast transformation protocol a previously described^44^. The primers used to amplify the fragments for assembly contained 20 bp binding and 40 bp of overlapping homology to the adjacent DNA fragment. A list of assembly primers for each plasmid is provided in Supplementary Table 3. Following assembly, DNA was isolated from the *S. cerevisiae* and the plasmid pool was electroporated into *E. coli* Epi300. Plasmids from individual colonies were screened for correct assembly using multiplex PCR and a diagnostic digest, and one final clone was confirmed by next-generation sequencing (MGH CCIB DNA Core, Massachusetts, USA).

All plasmids (pDEINO1-10) have a pCC1BAC-yeast backbone allowing replication and selection in *E. coli* and *S. cerevisiae* and have a low-copy *E. coli* origin that can be induced to high copy with arabinose. They also have an origin of transfer (*oriT*) necessary for direct DNA transfer via conjugation. Additional components on each plasmid are specified below.

The pDEINO1 replicating plasmid was constructed by restriction digestion of pAGE3.0^30^ with PacI to insert a synthesized codon-optimized *cat* gene under the control of a constitutive promoter (drKatA) and an origin of replication for *D. radiodurans* from pRadDEST-GFP and pRAD1^17^, respectively. This plasmid also contains elements for replication and selection in other alternative host organisms, including *S. meliloti* and *P. tricornutum*, and contains an *nptII* marker for *S. meliloti* that is functional in *D. radiodurans* as well.

The pDEINO2 nonreplicating plasmid contains a 1 kb region of homology to the *McrC* gene on the MP1 megaplasmid, amplified from wild type *D. radiodurans* genomic DNA and a codon-optimized *cat* gene for selection in *D. radiodurans*.

The pDEINO3-5 replicating plasmids contain an origin of replication for *D. radiodurans*, a codon-optimized *cat* gene, coupled with selective markers genes *tetR/A* from pAGE2.0, *aadA1* from pAGE1.0 and *aacC1* from pTA-Mob, respectively. The pDEINO6 plasmid has the same elements as pDEINO3, but without the codon-optimized *cat* gene.

The pDEINO7-10 nonreplicating plasmids contain two 1 kb regions of homology flanking ORF14075, ORF15360, *Mrr*, and ORF2230, respectively, amplified from wild type *D. radiodurans* genomic DNA. Between the homology regions on the plasmid is a selective marker, *tetR/A* for pDEINO8 and pDEINO9 or *nptII* for pDEINO7 and pDEINO10.

### Conjugation from *E. coli* to *D. radiodurans*

The *E. coli* ECGE101 *dapA* donor strain^30^, harboring pTA-Mob^36^ and pDEINO1, and a control strain, harboring only pDEINO1, were grown at 37°C overnight in 5 mL of LB media supplemented with diaminopimelic acid 60 μg mL^-1^, gentamicin 40 μg mL^-1^ (donor only), and chloramphenicol 15 μg^-1^. The saturated *E. coli* cultures were diluted 1:50 into 50 mL of the same media and grown for ∼2 h to an OD_600_ of 0.6. The *D. radiodurans* R1 recipient strain was grown at 30°C overnight in TGY media to an OD_600_ of 0.6. Donor, control, and recipient cultures were transferred to 50 mL falcon tubes and centrifuged at 3,000 x g for 20 min at 4°C. The supernatant was discarded, and cell pellets were resuspended in 300 μL of TGY media. Then, 50 μL of *E. coli* donor or control culture and 50 μL of *D. radiodurans* recipient culture were directly mixed and spread on a conjugation plate, previously dried for 1 h, consisting of TGY media supplemented with diaminopimelic acid 60 μg mL^-1^. After conjugation plates were incubated at 30°C for 3 h, the cells were scraped off with 1 mL of TGY media and adjusted to a final volume of 1 mL in a microfuge tube. Cell suspensions were serially diluted in TGY media and 100 μL was plated on both selective (TGY supplemented with chloramphenicol 5 μg mL^-1^) and nonselective (TGY) plates to select transconjugants and determine the conjugation frequency. Nonselective and selective plates were incubated at 30°C for 2 days and 3 days, respectively, and colonies were counted manually.

### Transconjugant Plasmid Isolation

*D. radiodurans* transconjugant colonies (10 from each experiment, 30 total) were passed twice on TGY plates supplemented with chloramphenicol (5 μg/mL), then inoculated into 5 mL of the same media and grown overnight. Alkaline lysis was performed using 3 mL of saturated culture as described^44^, and the DNA was electroporated into *E. coli* Epi300 cells. The plasmids were induced to high copy number in *E. coli* in 5 mL of LB media supplemented with chloramphenicol (15 μg/mL) and arabinose (100 μg/mL) for 8 hours before isolating for analysis using the BioBasic EZ10 Spin Column Miniprep Kit.

### Transconjugant Plasmid Analysis

Plasmids were analyzed by multiplex PCR using the primers listed in Supplementary Table 3 using 10 μL of Qiagen MPX, 3 μL of primer mix, 6 μL of water and 1 μL of template diluted to 1 ng μL^-1^. Thermocycler conditions were as follows: 95°C 5 min, 35 cycles of: 94°C 30 sec, 60°C 90 sec, and 72°C 10 sec, then 72°C 10 min. Gel electrophoresis was performed by loading 2 uL of the multiplex on a 2% agarose gel at 90 kV for 60 min and was stained with ethidium bromide for visualization. Transconjugant plasmids were further analyzed with a diagnostic restriction digest using 0.2 μL of EcoRI-HF, 5 μL of ∼100 ng/μL plasmid DNA, 2 μL Cutsmart buffer and 12.8 μL of water incubated at 37°C for 2 hours. Gel electrophoresis was performed by loading 10 uL of the digest on a 1% agarose gel at 100 kV for 120 min. Two transconjugant plasmids were analyzed by next-generation sequencing (MGH CCIB DNA Core, Massachusetts, USA) and compared to the sequence of the original pDEINO1 plasmid using the Benchling alignment tool (Benchling [Biology Software] (2021). Retrieved from https://benchling.com).

### Spot Plating *D. radiodurans*

*D. radiodurans* was grown overnight in 5 mL cultures of TGY media supplemented with the appropriate antibiotics for plasmid maintenance. The cultures were diluted to an OD_600_ of 0.1 before performing 10-fold serial dilutions in TGY media up to 10^−5^ dilution. Then, 5 uL of each dilution was plated on TGY plates supplemented with appropriate antibiotics and incubated at 30 degrees for 2-3 days.

### Plasmid Loss Stability Assay of *D. radiodurans* Transconjugants

One *D. radiodurans* transconjugant harboring pDEINO1 was plated on TGY supplemented with chloramphenicol (5 μg mL^-1^) to obtain single colonies. A single colony was inoculated in 50 mL of the same media and grown with shaking to an OD_600_ of 0.5 at 30°C. The culture was diluted to an OD_600_ of 0.1 and 100 uL of a dilution series of 10^−1^ to 10^−5^ was plated on non-selective (TGY) and selective (TGY supplemented with chloramphenicol 5 μg mL^-1^) media. These plates were incubated at 30°C for 3 days and colonies were counted manually. From the diluted culture (OD_600_ of 0.1), 50 uL was used to subculture into 50 mL of fresh non-selective media and grown with shaking at 30°C for an additional ∼13 generations to an OD_600_ of 0.5. This process was repeated for a total of ∼40 generations. The ratio of *D. radiodurans* colonies on selective plates to colonies on non-selective plates after each subculturing event was reported as an indicator of plasmid loss. Additionally, 100 colonies from the selective and nonselective plates were struck onto TGY plates supplemented with chloramphenicol each day and the number of colonies unable to grow on selection was used as a second strategy to determine plasmid loss.

### Cloning the *D. radiodurans* MP1 Megaplasmid

Conjugation from *E. coli* to *D. radiodurans* was performed as described above using *E. coli* ECGE101 pTA-Mob pDEINO2 as the donor and *D. radiodurans* as the recipient, with *D. radiodurans* grown to an OD_600_ of 2 rather than 0.6. *D. radiodurans* transconjugants were selected by plating 100 μL onto 10 TGY plates supplemented with chloramphenicol (5 μg/mL). Three transconjugants were isolated and analyzed as previously described, transformed into *E. coli* Epi300 and, following induction in *E. coli*, plasmids were isolated using alkaline lysis. The cloned MP1 plasmids were analyzed by two multiplex PCRs, one amplifying fragments within the conjugative suicide plasmid, and one amplifying fragments within the MP1 megaplasmid. The plasmids were further analyzed by two diagnostic restriction digests using MfeI-HF and NheI-HF. Finally, total DNA from one clone was isolated using the Monarch Kit for HMW DNA Extraction from Bacteria (NEB#T3060) and confirmed by Nanopore sequencing.

### Gene Deletions

Conjugation from *E. coli* to *D. radiodurans* was performed as described above using *E. coli* ECGE101 pTA-Mob harboring pDEINO7 (ORF14075), pDEINO8 (ORF15360), pDEINO9 (*Mrr*) or pDEINO10 (ORF2230) as the donor and *D. radiodurans* as the recipient. *D. radiodurans* transconjugants were selected on TGY supplemented with neomycin (5 μg/mL) for pDEINO7 and pDEINO10 or tetracycline (0.5 μg/mL) for pDEINO8 and pDEINO9. Transconjugants were isolated and analyzed by multiplex PCR with primers listed in Supplementary Table 3 using 10 μL of Qiagen MPX, 2 μL of primer mix, 6 μL of water, 1 μL of DMSO and 1 μL of alkaline lysis DNA. Thermocycler conditions were as follows: 95°C 5 min, 30 cycles of 94°C 30 sec, 60°C 90 sec, and 72°C 90 sec, then 72°C 10 min. Gel electrophoresis was performed by loading 2 uL of the multiplex on a 2% agarose gel at 110 kV for 50 min.

### *D. radiodurans* R1 Genomic DNA Isolation

*D. radiodurans* genomic DNA was isolated in agar plugs using the Bio-Rad CHEF Genomic DNA Plug Kit (Bio-Rad, CAT#170-3592) with an adapted protocol^44^. To prepare the plugs, 50 mL of *D. radiodurans* culture was grown to OD600 of 1.0, chloramphenicol (100 μg mL-1) was added and the culture was grown for an additional hour. The culture was centrifuged at 5000 × g for 5 min at 4°C. Cells were washed once with 1 M sorbitol in 1.5 mL Eppendorf tubes and centrifuged at 4000 RPM for 3 min. The supernatant was removed, and cells were resus-pended in 600 uL of protoplasting solution (for 10 mL: 4.56 mL of SPEM solution, 1000 μl Zymolyase-20 T solution (50 mg mL^-1^ dissolved in H_2_O), 400 μL lysozyme (25 mg mL^-1^), 400 μl hemicellulase (25 mg mL^-1^), 50 μl β-Mercaptoethanol). The cell suspension was incubated for 5 min at 37°C and mixed with an equal volume of 2.0% low melting-point agarose in 1 × TAE buffer (40 mM Tris, 20 mM acetic acid and 1 mM EDTA) which was equilibrated at 50°C. Aliquots of 95 μl were transferred into plug molds (Bio-Rad, CAT#170– 3713) and allowed to solidify for 10 min at 4°C. Next, plugs were removed from the molds into 50 ml conical tube containing 5 mL of protoplasting solution and incubated for 45 min at 37°C. Next, plugs were washed with 25 ml of wash buffer (20 mM Tris, 50 mM EDTA, pH 8.0), and then incubated in 5 ml in Proteinase K buffer (100 mM EDTA (pH 8.0), 0.2% sodium deoxycholate, and 1% sodium lauryl sarcosine, 1 mg ml-1 Proteinase K) for 24 hrs at 50°C. The plugs were washed 4 times with 25 mL of wash buffer for 30 min each at room temperature and incubated in wash buffer overnight at 4°C. The next day, the plugs were washed 4 times with 10X diluted wash buffer for 30 minutes each, then stored at 4°C in 10X diluted wash buffer. To isolate DNA from the plugs, two plugs were transferred to a 1.5 mL Eppendorf tube and were washed once with 10X diluted wash buffer for 1 hour, and once with TE buffer for 1 hour. The TE buffer was removed and the tube was incubated in a 42°C water bath for 10 min, followed by a 65°C water bath for 10 min. The tube was returned to the 42°C water bath for 10 min then 50 μL of TE buffer and 3 μL of β-agarase was added and flicked gently to mix. The plugs were incubated at 42°C for an hour. Another 50 μL of TE buffer was added to the tube and left for 2 hours at 42°C. The DNA was checked for purity by gel electrophoresis of 1 μL on a 1% gel, run at 100 kV for 40 min.

### DNA Sequencing and Analysis

*D. radiodurans R1*. The library was prepared using the SQK-LSK109 kit according to the manufacturer’s protocol. An R9.4.1 flow cell was used for sequencing. Basecalling was performed using Guppy v4.2.2 (Oxford Nanopore Technologies) in high-accuracy mode.

Genome assembly was performed with Flye v2.8.1^48^. The assembly was polished with one round of Racon^49^ and one round of Medaka (Oxford Nanopore Technologies). *E. coli pDEINO2-MP1*. The library was prepared using the SQK-LSK109 kit and the EXP-NBD104 native barcoding kit according to the manufacturer’s protocol. An R9.4.1 flow cell was used for sequencing. Basecalling was performed using Guppy v5.0.7 (Oxford Nanopore Technologies) in high-accuracy mode. Genome assembly was performed with Flye v2.8.1 in –meta mode^48^. The assembly was polished with one round of Racon^49^ and one round of Medaka (Oxford Nanopore Technologies). Plasmids were extracted from the final assemblies, and aligned using minimap2^50^ against the expected sequences to identify any mutations.

## ASSOCIATED CONTENT

### Supporting Information

Supplementary File

## AUTHOR INFORMATION

### Author Contributions

S.L.B., D.R.E and B.J.K conceived the experiments, S.L.B, K.D.V.B and D.J.G. conducted the experiments, S.L.B, D.J.G and B.J.K analyzed the results, S.L.B. and B.J.K wrote the paper. All authors edited the paper. All authors have given approval to the final version of the manuscript.

### Notes

The authors declare no competing financial interests.

## ACKNOWLEDGMENT

This research was funded by Natural Sciences and Engineering Research Council of Canada (NSERC), grant number: RGPIN-2018-06172. The authors would like to thank Dr. Murray Junop and Robert Szabla (Western University) for providing the *D. radiodurans* R1 strain, pRAD1 and pRadDEST-GFP plasmids. We also wish to thank Dr. Gregory Gloor for providing the resources needed for Oxford Nanopore sequencing and genome assembly. In addition to the figures noted, the graphical abstract was also created with BioRender.com.

