## Supplemental file for "Conjugation-based genome engineering in *Deinococcus radiodurans*"

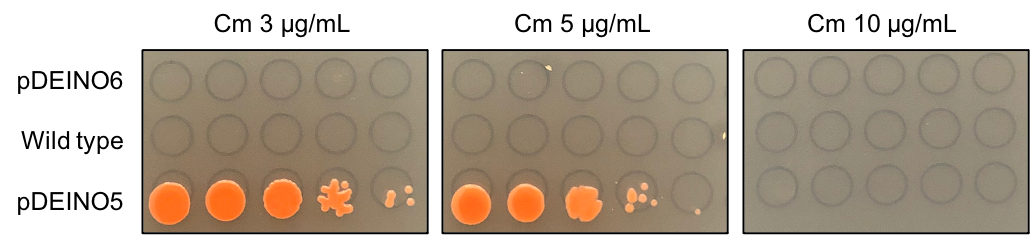

**Supplementary Figure 1.** Test of chloramphenicol resistance in *D. radiodurans* harboring pDEINO6 (containing *E. coli* *cat* gene only) and pDEINO5 (containing *D. radiodurans* codon optimized *cat* gene and *E. coli* *cat* gene) compared to wild type. A 10-fold dilution series of each strain was spot plated on TGY media supplemented with increasing concentration of chloramphenicol.

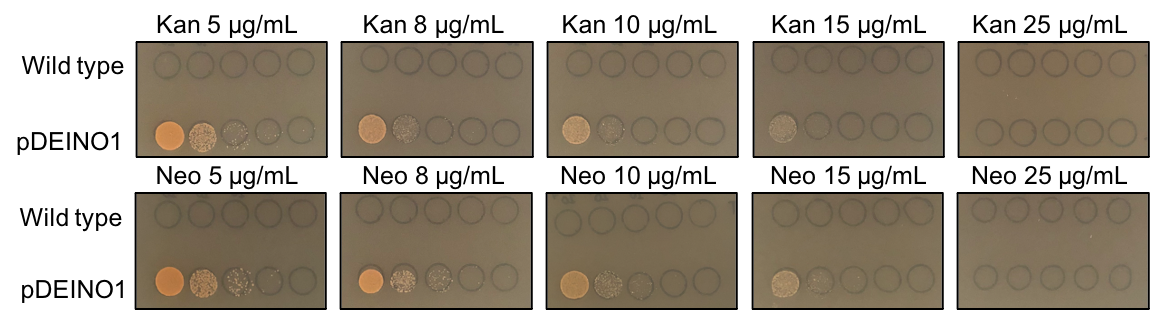

**Supplementary Figure 2.** Test of kanamycin and neomycin sensitivity of *D. radiodurans* harboring pDEINO1 compared to wild type. A 10-fold dilution series of each strain was spot plated on TGY media supplemented with increasing concentration of antibiotic.

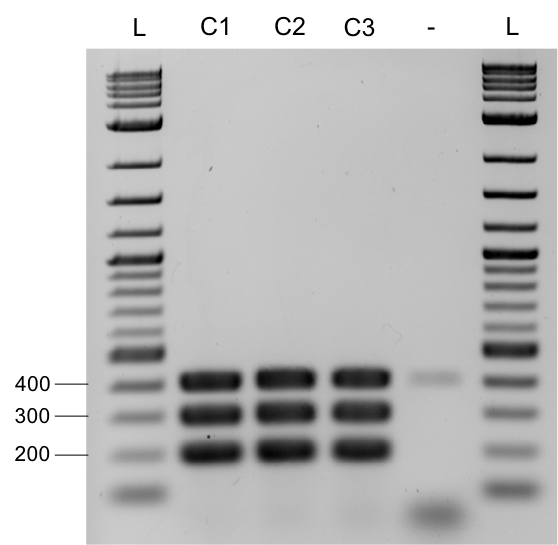

**Supplementary Figure 3.** Agarose gel of multiplex PCR performed on DNA extracted from three *D. radiodurans* transconjugant colonies following conjugation of pDEINO2 to clone the MP1 megaplasmid. Amplicons from the integrated pDEINO2 plasmid should be 200, 300 and 400 bp in size. L, 2-log ladder. -, water.

**Supplementary Table 1.** Plasmid-loss stability assay of *D. radiodurans* harboring pDEINO1 over 40 generations**.** The nonselective media was TGY, while selective media was TGY Cm 5 μg mL^-1^.

| **Day Number** | **Colonies on Nonselective Media (x 10^4^)** | **Colonies on Selective Media (x 10^3^)** | **Surviving Colonies Restruck on Selective Media** |
| --- | --- | --- | --- |
| 1 | 116 | 282 | 71 |
| 2 | 105 | 109 | 32 |
| 3 | 140 | 55 | 14 |
| 4 | 106 | 11 | 6 |

**Supplementary Table 2.** List of plasmids used in this study.

| **Plasmid** | **Description** | **Resistance** | **Reference or Source** |
| --- | --- | --- | --- |
| pAGE1.0 | MHS vector | Cm (*E. coli*), HIS3 (*S. cerevisiae*),  Str/Spec (*S. meliloti*), Ntc (*P. tricornutum*) | [1] |
| pAGE2.0 | MHS vector | Cm (*E. coli*), HIS3 (*S. cerevisiae*),  Tet (*S. meliloti*), Ntc (*P. tricornutum*) | [1] |
| pAGE3.0 | MHS vector | Cm (*E. coli*), HIS3 (*S. cerevisiae*),  Nm (*S. meliloti*), Ntc (*P. tricornutum*) | [1] |
| pRAD1 | General cloning vector for use in *E. coli* or *D. radiodurans* | Amp (*E. coli),* Cm (*D. radiodurans*) | [2] |
| pTA-Mob | Broad-host-range mobilization plasmid | Gm (*E. coli)* | [3] |
| pRadDEST-GFP |  |  | Unpublished |
| pDEINO1 | pAGE3.0 with *D. radiodurans* origin and codon-optimized Cm marker | Cm (*D. radiodurans*), Cm (*E. coli*), HIS3 (*S. cerevisiae*), Nm (*S. meliloti* and *D. radiodurans*), Ntc (*P. tricornutum*) | This study |
| pDEINO2 | Nonreplicating plasmid with 1 kb homology to *McrC* | Cm (*D. radiodurans*), Cm (*E. coli*), HIS3 (*S. cerevisiae*) | This study |
| pDEINO3 | Replicating plasmid with Tet selection marker | Tet and Cm (*D. radiodurans*),  Cm (*E. coli*), HIS3 (*S. cerevisiae*) | This study |
| pDEINO4 | Replicating plasmid with Str selection marker | Str and Cm (*D. radiodurans*),  Cm (*E. coli*), HIS3 (*S. cerevisiae*) | This study |
| pDEINO5 | Replicating plasmid with Gm marker | Gm and Cm (*D. radiodurans*),  Cm (*E. coli*), HIS3 (*S. cerevisiae*) | This study |
| pDEINO6 | Replicating plasmid with Tet selection marker and without *D. radiodurans* Cm | Tet (*D. radiodurans*), Cm (*E. coli*), HIS3 (*S. cerevisiae*) | This study |
| pDEINO7 | Nonreplicating plasmid with two 1 kb homology regions flanking ORF14075 | Nm (*D. radiodurans*), Cm (*E. coli*), HIS3 (*S. cerevisiae*) | This study |
| pDEINO8 | Nonreplicating plasmid with two 1 kb homology regions flanking ORF15360 | Tet (*D. radiodurans*), Cm (*E. coli*), HIS3 (*S. cerevisiae*) | This study |
| pDEINO9 | Nonreplicating plasmid with two 1 kb homology regions flanking *Mrr* | Tet (*D. radiodurans*), Cm (*E. coli*), HIS3 (*S. cerevisiae*) | This study |
| pDEINO10 | Nonreplicating plasmid with two 1 kb homology regions flanking ORF2230 | Nm (*D. radiodurans*), Cm (*E. coli*), HIS3 (*S. cerevisiae*) | This study |

**Supplementary Table 3.** List of oligonucleotides used in this study. The bold, underlined sequence in the assembly primers represents the binding portion of the primer, while the remainder of the sequence is the hook or homology region to the adjacent fragment.

| **Name** | **Sequence (5’ to 3’)** | **Description** |
| --- | --- | --- |
| pDEINO1 Assembly Primers | | |
| BK486_F | GTTCCGCTTCCTTTAGCAGCCCTTGCGCCCTGAGTGCTTGCGGCAGCGTGAAGCTTTAAT**CATGATTACGCCAAGCTCGC** | pDEINO1 assembly primer  (origin from pRAD1) |
| BK486_R | ATATTGAGAATCATTCTCAATGTCCAGGGCCCTCGGTCTCCATGGCCCTCAGGCCCTCGC**TTTAGCTTCCTTAGCTCCTG** | pDEINO1 assembly primer  (origin from pRAD1) |
| BK487_F | GCTTGAGTAGGACAAATCCGCCGAGCTTCGACGAGATTTTCAGGAGCTAAGGAAGCTAAA**GCGAGGGCCTGAGGGCCATG** | pDEINO1 assembly primer  (DrCm^R^) |
| BK487_R | GGATATACCGAAAAAATCGCTATAATGACCCCGAAGCAGGGTTATGCAGCGGAAGATTTA**AAAAAACCCCCCGGATTGCC** | pDEINO1 assembly primer  (DrCm^R^) |
| pDEINO2 Assembly Primers | | |
| BK1388_F | AGTACATCACCGACGAGCAAGGCAAGACGATCTTAATTAA**TCGAGCTGGTTGCCCTCGCC** | pDEINO2 assembly primer (split Pcc1BAC-yeast #1) |
| BK1388_R | GAAGAGCGTTGATCAATGGCCTGTTCAAAAACAGTTCTCA**TCCGGATCTGACCTTTACCA** | pDEINO2 assembly primer (split Pcc1BAC-yeast #1) |
| BK1389_F | GTACGTGAAACGGATGAAGTTGGTAAAGGTCAGATCCGGA**TGAGAACTGTTTTTGAACAG** | pDEINO2 assembly primer (split Pcc1BAC-yeast #2) |
| BK1409_R | CAAATGCTCGCCTATAGCGAGGCCTTTCAGCACAGCGCGG**GTTTAAACGGGCTTCGCCCT** | pDEINO2 assembly primer (split Pcc1BAC-yeast #2) |
| BK1410_F | GCTCGCCGCAGTCGAGCGACAGGGCGAAGCCCGTTTAAAC**CCGCGCTGTGCTGAAAGGCC** | pDEINO2 assembly primer (1 kb homology to DraR1McrCP) |
| BK1410_R | GCCCTCGGTCTCCATGGCCCTCAGGCCCTCGCGGCGCGCC**CTGAGCCCGGTGGCCACCAT** | pDEINO2 assembly primer (1 kb homology to DraR1McrCP) |
| BK1411_F | TCAAAGCGGGTCATGGTGGCCACCGGGCTCAGGGCGCGCC**GCGAGGGCCTGAGGGCCATG** | pDEINO2 assembly primer (DrCm^R^ and oriT) |
| BK1392_R | CGCCAGCCCAGCGGCGAGGGCAACCAGCTCGATTAATTAA**GATCGTCTTGCCTTGCTCGT** | pDEINO2 assembly primer (DrCm^R^ and oriT) |
| pDEINO20 Assembly Primers | | |
| BK1388_F | AGTACATCACCGACGAGCAAGGCAAGACGATCTTAATTAA**TCGAGCTGGTTGCCCTCGCC** | pDEINO20 assembly primer (split Pcc1BAC-yeast #1) |
| BK1388_R | GAAGAGCGTTGATCAATGGCCTGTTCAAAAACAGTTCTCA**TCCGGATCTGACCTTTACCA** | pDEINO20 assembly primer (split Pcc1BAC-yeast #1) |
| BK1389_F | GTACGTGAAACGGATGAAGTTGGTAAAGGTCAGATCCGGA**TGAGAACTGTTTTTGAACAG** | pDEINO20 assembly primer (split Pcc1BAC-yeast #2) |
| BK1389_R | TCGATAGATCTCGAGGCCTCGCGAGCTTGGCGTAATCATG**GTTTAAACGGGCTTCGCCCT** | pDEINO20 assembly primer (split Pcc1BAC-yeast #2) |
| BK1390_F | GCTCGCCGCAGTCGAGCGACAGGGCGAAGCCCGTTTAAAC**CATGATTACGCCAAGCTCGC** | pDEINO20 assembly primer (Drad origin) |
| BK1694_R | CACGGCCGCGCTCGGCCTCTCTGGCGGCCTTCTGGCGCTC**TTTAGCTTCCTTAGCTCCTG** | pDEINO20 assembly primer (Drad origin) |
| BK1695_F | CCGAGCTTCGACGAGATTTTCAGGAGCTAAGGAAGCTAAA**GAGCGCCAGAAGGCCGCCAG** | pDEINO20 assembly primer (Tet^R^) |
| BK1695_R | TGTCCAGGGCCCTCGGTCTCCATGGCCCTCAGGCCCTCGC**GATCAGACGCTGAGTGCGCT** | pDEINO20 assembly primer (Tet^R^) |
| BK1696_F | GCAGGACGCCGATGATTTGAAGCGCACTCAGCGTCTGATC**GCGAGGGCCTGAGGGCCATG** | pDEINO20 assembly primer (DrCm^R^ and oriT) |
| BK1392_R | CGCCAGCCCAGCGGCGAGGGCAACCAGCTCGATTAATTAA**GATCGTCTTGCCTTGCTCGT** | pDEINO20 assembly primer (DrCm^R^ and oriT) |
| pDEINO29 Assembly Primers | | |
| BK1388_F | AGTACATCACCGACGAGCAAGGCAAGACGATCTTAATTAA**TCGAGCTGGTTGCCCTCGCC** | pDEINO29 assembly primer (split Pcc1BAC-yeast #1) |
| BK1388_R | GAAGAGCGTTGATCAATGGCCTGTTCAAAAACAGTTCTCA**TCCGGATCTGACCTTTACCA** | pDEINO29 assembly primer (split Pcc1BAC-yeast #1) |
| BK1389_F | GTACGTGAAACGGATGAAGTTGGTAAAGGTCAGATCCGGA**TGAGAACTGTTTTTGAACAG** | pDEINO29 assembly primer (split Pcc1BAC-yeast #2) |
| BK1874_R | ACCTTGCGGTACCGGGCGAGGCATTACCCTGTTATCCCTA**GGGCTTCGCCCTGTCGCTCG** | pDEINO29 assembly primer (split Pcc1BAC-yeast #2) |
| BK1875_F | GTCGAGCGACAGGGCGAAGCCCTAGGGATAACAGGGTAAT**GCCTCGCCCGGTACCGCAAG** | pDEINO29 assembly primer (ORF14075 homology #1) |
| BK1875_R | CCAGCGCGGGGATCTCATGCTGGAGTTCTTCGCCCACCCC**ATCAGCTCGCAGGCTAGCGC** | pDEINO29 assembly primer (ORF14075 homology #1) |
| BK1876_F | CTCCGACTGACTTTGCTGCTGCGCTAGCCTGCGAGCTGAT**GGGGTGGGCGAAGAACTCCA** | pDEINO29 assembly primer (NptII^R^) |
| BK1876_R | CAAGAAAGCCTCGTCGTAAGGCGAGTACGAATCTCAAGCT**AGCTTCACGCTGCCGCAAGC** | pDEINO29 assembly primer (NptII^R^) |
| BK1877_F | GCAGCCCTTGCGCCCTGAGTGCTTGCGGCAGCGTGAAGCT**AGCTTGAGATTCGTACTCGC** | pDEINO29 assembly primer (ORF14075 homology #2) |
| BK1877_R | GCAGGGTTATGCAGCGGAAGATATTACCCTGTTATCCCTA**CCCAGACCTGCTCCGGCGTG** | pDEINO29 assembly primer (ORF14075 homology #2) |
| BK1878_F | GGCACGCCGGAGCAGGTCTGGGTAGGGATAACAGGGTAAT**ATCTTCCGCTGCATAACCCT** | pDEINO29 assembly primer (oriT) |
| BK1392_R | CGCCAGCCCAGCGGCGAGGGCAACCAGCTCGATTAATTAA**GATCGTCTTGCCTTGCTCGT** | pDEINO29 assembly primer (oriT) |
| pDEINO30 Assembly Primers | | |
| BK1388_F | AGTACATCACCGACGAGCAAGGCAAGACGATCTTAATTAA**TCGAGCTGGTTGCCCTCGCC** | pDEINO30 assembly primer (split Pcc1BAC-yeast #1) |
| BK1388_R | GAAGAGCGTTGATCAATGGCCTGTTCAAAAACAGTTCTCA**TCCGGATCTGACCTTTACCA** | pDEINO30 assembly primer (split Pcc1BAC-yeast #1) |
| BK1389_F | GTACGTGAAACGGATGAAGTTGGTAAAGGTCAGATCCGGA**TGAGAACTGTTTTTGAACAG** | pDEINO30 assembly primer (split Pcc1BAC-yeast #2) |
| BK1879_R | CAGTCCTCGGAAACTTCTGCCCATTACCCTGTTATCCCTA**GGGCTTCGCCCTGTCGCTCG** | pDEINO30 assembly primer (split Pcc1BAC-yeast #2) |
| BK1880_F | GTCGAGCGACAGGGCGAAGCCCTAGGGATAACAGGGTAAT**GGGCAGAAGTTTCCGAGGAC** | pDEINO30 assembly primer (ORF15360 homology #1) |
| BK1880_R | CACGGCCGCGCTCGGCCTCTCTGGCGGCCTTCTGGCGCTC**ACAATGCCATTTATGTTTTC** | pDEINO30 assembly primer (ORF15360 homology #1) |
| BK1881_F | CAGCTAGTGCCGAAGTGCCAGAAAACATAAATGGCATTGT**GAGCGCCAGAAGGCCGCCAG** | pDEINO30 assembly primer (Tet^R^) |
| BK1881_R | AACTTTCCGACCGCGCCCAGACGGGCAAGGTCGAGGGGGC**GATCAGACGCTGAGTGCGCT** | pDEINO30 assembly primer (Tet^R^) |
| BK1882_F | GCAGGACGCCGATGATTTGAAGCGCACTCAGCGTCTGATC**GCCCCCTCGACCTTGCCCGT** | pDEINO30 assembly primer (ORF15360 homology #2) |
| BK1882_R | GCAGGGTTATGCAGCGGAAGATATTACCCTGTTATCCCTA**GTGATTCAGCCGCTGCTCTT** | pDEINO30 assembly primer (ORF15360 homology #2) |
| BK1883_F | CAAAGAGCAGCGGCTGAATCACTAGGGATAACAGGGTAAT**ATCTTCCGCTGCATAACCCT** | pDEINO30 assembly primer (oriT) |
| BK1392_R | CGCCAGCCCAGCGGCGAGGGCAACCAGCTCGATTAATTAA**GATCGTCTTGCCTTGCTCGT** | pDEINO30 assembly primer (oriT) |
| pDEINO32 Assembly Primers | | |
| BK1388_F | AGTACATCACCGACGAGCAAGGCAAGACGATCTTAATTAA**TCGAGCTGGTTGCCCTCGCC** | pDEINO32 assembly primer (split Pcc1BAC-yeast #1) |
| BK1388_R | GAAGAGCGTTGATCAATGGCCTGTTCAAAAACAGTTCTCA**TCCGGATCTGACCTTTACCA** | pDEINO32 assembly primer (split Pcc1BAC-yeast #1) |
| BK1389_F | GTACGTGAAACGGATGAAGTTGGTAAAGGTCAGATCCGGA**TGAGAACTGTTTTTGAACAG** | pDEINO32 assembly primer (split Pcc1BAC-yeast #2) |
| BK1869_R | GTACGAGCAGGAACTGGGATTCATTACCCTGTTATCCCTA**GGGCTTCGCCCTGTCGCTCG** | pDEINO32 assembly primer (split Pcc1BAC-yeast #2) |
| BK1870_F | GTCGAGCGACAGGGCGAAGCCCTAGGGATAACAGGGTAAT**GAATCCCAGTTCCTGCTCGT** | pDEINO32 assembly primer (Mrr homology #1) |
| BK2124_R | CACGGCCGCGCTCGGCCTCTCTGGCGGCCTTCTGGCGCTC**ATTTTTTAAGTTTACGCTCT** | pDEINO32 assembly primer (Mrr homology #1) |
| BK2125_F | CGAAAGCCGTAAGTCCAGACAGAGCGTAAACTTAAAAAAT**GAGCGCCAGAAGGCCGCCAG** | pDEINO32 assembly primer (Tet^R^) |
| BK2125_R | CCTCCTCCCTTGACGCCGGAAACCGCCCTGTCAGGCGAGC**GATCAGACGCTGAGTGCGCT** | pDEINO32 assembly primer (Tet^R^) |
| BK2126_F | GCAGGACGCCGATGATTTGAAGCGCACTCAGCGTCTGATC**GCTCGCCTGACAGGGCGGTT** | pDEINO32 assembly primer (Mrr homology #2) |
| BK1872_R | GCAGGGTTATGCAGCGGAAGATATTACCCTGTTATCCCTA**CGGCAGTTCCACGTTGCACA** | pDEINO32 assembly primer (Mrr homology #2) |
| BK1873_F | GTTGTGCAACGTGGAACTGCCGTAGGGATAACAGGGTAAT**ATCTTCCGCTGCATAACCCT** | pDEINO32 assembly primer (oriT) |
| BK1392_R | CGCCAGCCCAGCGGCGAGGGCAACCAGCTCGATTAATTAA**GATCGTCTTGCCTTGCTCGT** | pDEINO32 assembly primer (oriT) |
| pDEINO33 Assembly Primers | | |
| BK1388_F | AGTACATCACCGACGAGCAAGGCAAGACGATCTTAATTAA**TCGAGCTGGTTGCCCTCGCC** | pDEINO33 assembly primer (split Pcc1BAC-yeast #1) |
| BK1388_R | GAAGAGCGTTGATCAATGGCCTGTTCAAAAACAGTTCTCA**TCCGGATCTGACCTTTACCA** | pDEINO33 assembly primer (split Pcc1BAC-yeast #1) |
| BK1389_F | GTACGTGAAACGGATGAAGTTGGTAAAGGTCAGATCCGGA**TGAGAACTGTTTTTGAACAG** | pDEINO33 assembly primer (split Pcc1BAC-yeast #2) |
| BK1864_R | GCGAGCATGTACGCCTGGGCGCATTACCCTGTTATCCCTA**GGGCTTCGCCCTGTCGCTCG** | pDEINO33 assembly primer (split Pcc1BAC-yeast #2) |
| BK1865_F | GTCGAGCGACAGGGCGAAGCCCTAGGGATAACAGGGTAAT**GCGCCCAGGCGTACATGCTC** | pDEINO33 assembly primer (ORF2230 homology #1) |
| BK2127_R | CCAGCGCGGGGATCTCATGCTGGAGTTCTTCGCCCACCCC**GAACCTCTTCAGAGTACGGC** | pDEINO33 assembly primer (ORF2230 homology #1) |
| BK2128_F | CCCACATGGCCCGAGTGTAAGCCGTACTCTGAAGAGGTTC**GGGGTGGGCGAAGAACTCCA** | pDEINO33 assembly primer (NptII^R^) |
| BK2128_R | CGGGCACCCGGGAGCTTCCCTGGGTGCCCGCTTTCGGTGT**AGCTTCACGCTGCCGCAAGC** | pDEINO33 assembly primer (NptII^R^) |
| BK2129_F | GCAGCCCTTGCGCCCTGAGTGCTTGCGGCAGCGTGAAGCT**ACACCGAAAGCGGGCACCCA** | pDEINO33 assembly primer (ORF2230 homology #2) |
| BK1867_R | GCAGGGTTATGCAGCGGAAGATATTACCCTGTTATCCCTA**CGTGAGCACCATTTTCATCC** | pDEINO33 assembly primer (ORF2230 homology #2) |
| BK1868_F | CAGGATGAAAATGGTGCTCACGTAGGGATAACAGGGTAAT**ATCTTCCGCTGCATAACCCT** | pDEINO33 assembly primer (oriT) |
| BK1392_R | CGCCAGCCCAGCGGCGAGGGCAACCAGCTCGATTAATTAA**GATCGTCTTGCCTTGCTCGT** | pDEINO33 assembly primer (oriT) |
| pDEINO37 Assembly Primers | | |
| BK1388_F | AGTACATCACCGACGAGCAAGGCAAGACGATCTTAATTAA**TCGAGCTGGTTGCCCTCGCC** | pDEINO37 assembly primer (split Pcc1BAC-yeast #1) |
| BK1388_R | GAAGAGCGTTGATCAATGGCCTGTTCAAAAACAGTTCTCA**TCCGGATCTGACCTTTACCA** | pDEINO37 assembly primer (split Pcc1BAC-yeast #1) |
| BK1389_F | GTACGTGAAACGGATGAAGTTGGTAAAGGTCAGATCCGGA**TGAGAACTGTTTTTGAACAG** | pDEINO37 assembly primer (split Pcc1BAC-yeast #2) |
| BK1389_R | TCGATAGATCTCGAGGCCTCGCGAGCTTGGCGTAATCATG**GTTTAAACGGGCTTCGCCCT** | pDEINO37 assembly primer (split Pcc1BAC-yeast #2) |
| BK1390_F | GCTCGCCGCAGTCGAGCGACAGGGCGAAGCCCGTTTAAAC**CATGATTACGCCAAGCTCGC** | pDEINO37 assembly primer (Drad origin) |
| BK1697_R | ACAAGCATAAAGCTTGCTCAATCAATCACCGGATCCCCGG**TTTAGCTTCCTTAGCTCCTG** | pDEINO37 assembly primer (Drad origin) |
| BK1698_F | CCGAGCTTCGACGAGATTTTCAGGAGCTAAGGAAGCTAAA**CCGGGGATCCGGTGATTGAT** | pDEINO37 assembly primer (Str^R^) |
| BK1698_R | TGTCCAGGGCCCTCGGTCTCCATGGCCCTCAGGCCCTCGC**GGATCCGGTGATTGATTGAG** | pDEINO37 assembly primer (Str^R^) |
| BK1699_F | GTTTACAAGCATAAAGCTTGCTCAATCAATCACCGGATCC**GCGAGGGCCTGAGGGCCATG** | pDEINO37 assembly primer (DrCm^R^ and oriT) |
| BK1392_R | CGCCAGCCCAGCGGCGAGGGCAACCAGCTCGATTAATTAA**GATCGTCTTGCCTTGCTCGT** | pDEINO37 assembly primer (DrCm^R^ and oriT) |
| pDEINO39 Assembly Primers | | |
| BK1388_F | AGTACATCACCGACGAGCAAGGCAAGACGATCTTAATTAA**TCGAGCTGGTTGCCCTCGCC** | pDEINO39 assembly primer (split Pcc1BAC-yeast #1) |
| BK1388_R | GAAGAGCGTTGATCAATGGCCTGTTCAAAAACAGTTCTCA**TCCGGATCTGACCTTTACCA** | pDEINO39 assembly primer (split Pcc1BAC-yeast #1) |
| BK1389_F | GTACGTGAAACGGATGAAGTTGGTAAAGGTCAGATCCGGA**TGAGAACTGTTTTTGAACAG** | pDEINO39 assembly primer (split Pcc1BAC-yeast #2) |
| BK1389_R | TCGATAGATCTCGAGGCCTCGCGAGCTTGGCGTAATCATG**GTTTAAACGGGCTTCGCCCT** | pDEINO39 assembly primer (split Pcc1BAC-yeast #2) |
| BK1390_F | GCTCGCCGCAGTCGAGCGACAGGGCGAAGCCCGTTTAAAC**CATGATTACGCCAAGCTCGC** | pDEINO39 assembly primer (Drad origin) |
| BK2117_R | CCCGGCCGCGGAGTTGTTCGGTAAATTGTCACAACGCCGC**TTTAGCTTCCTTAGCTCCTG** | pDEINO39 assembly primer (Drad origin) |
| BK2118_F | CCGAGCTTCGACGAGATTTTCAGGAGCTAAGGAAGCTAAA**GCGGCGTTGTGACAATTTAC** | pDEINO39 assembly primer (Gm^R^) |
| BK2118_R | TGTCCAGGGCCCTCGGTCTCCATGGCCCTCAGGCCCTCGC**GACGCACACCGTGGAAACGG** | pDEINO39 assembly primer (Gm^R^) |
| BK2119_F | AACTGGGTTCGTGCCTTCATCCGTTTCCACGGTGTGCGTC**GCGAGGGCCTGAGGGCCATG** | pDEINO39 assembly primer (DrCm^R^ and oriT) |
| BK1392_R | CGCCAGCCCAGCGGCGAGGGCAACCAGCTCGATTAATTAA**GATCGTCTTGCCTTGCTCGT** | pDEINO39 assembly primer (DrCm^R^ and oriT) |
| pDEINO40 Assembly Primers | | |
| BK1388_F | AGTACATCACCGACGAGCAAGGCAAGACGATCTTAATTAA**TCGAGCTGGTTGCCCTCGCC** | pDEINO40 assembly primer (split Pcc1BAC-yeast #1) |
| BK1388_R | GAAGAGCGTTGATCAATGGCCTGTTCAAAAACAGTTCTCA**TCCGGATCTGACCTTTACCA** | pDEINO40 assembly primer (split Pcc1BAC-yeast #1) |
| BK1389_F | GTACGTGAAACGGATGAAGTTGGTAAAGGTCAGATCCGGA**TGAGAACTGTTTTTGAACAG** | pDEINO40 assembly primer (split Pcc1BAC-yeast #2) |
| BK1389_R | TCGATAGATCTCGAGGCCTCGCGAGCTTGGCGTAATCATG**GTTTAAACGGGCTTCGCCCT** | pDEINO40 assembly primer (split Pcc1BAC-yeast #2) |
| BK1390_F | GCTCGCCGCAGTCGAGCGACAGGGCGAAGCCCGTTTAAAC**CATGATTACGCCAAGCTCGC** | pDEINO40 assembly primer (Drad origin) |
| BK1694_R | CACGGCCGCGCTCGGCCTCTCTGGCGGCCTTCTGGCGCTC**TTTAGCTTCCTTAGCTCCTG** | pDEINO40 assembly primer (Drad origin) |
| BK1695_F | CCGAGCTTCGACGAGATTTTCAGGAGCTAAGGAAGCTAAA**GAGCGCCAGAAGGCCGCCAG** | pDEINO40 assembly primer (Tet^R^) |
| BK2103_R | CGCTATAATGACCCCGAAGCAGGGTTATGCAGCGGAAGAT**GATCAGACGCTGAGTGCGCT** | pDEINO40 assembly primer (Tet^R^) |
| BK2104_F | GCAGGACGCCGATGATTTGAAGCGCACTCAGCGTCTGATC**ATCTTCCGCTGCATAACCCT** | pDEINO40 assembly primer (oriT) |
| BK1392_R | CGCCAGCCCAGCGGCGAGGGCAACCAGCTCGATTAATTAA**GATCGTCTTGCCTTGCTCGT** | pDEINO40 assembly primer (oriT) |
| pDEINO1 MPX Primers | | |
| BK1327_F | GTGGAGCGGATTATGTCAGCAATG | pDEINO1 MPX primers – 200 bp (junction between pCC1BAC-yeast and NptII) |
| BK1327_R | CGGAATCGTTTTCCGGGACG | pDEINO1 MPX primers – 200 bp (junction between pCC1BAC-yeast and NptII) |
| BK1328_F | CCGCAATACCGGCTTCGCT | pDEINO1 MPX primers – 300 bp (within RepA2B2C2) |
| BK1328_R | CTCGAGGGCCTCTTCTGGAAGG | pDEINO1 MPX primers – 300 bp (within RepA2B2C2) |
| BK1329_F | GCAGTAGCAGAACAGGCCACAC | pDEINO1 MPX primers – 400 bp (within pCC1BAC-yeast) |
| BK1329_R | GGGTAATTCTGCTAGCCTCTGCAA | pDEINO1 MPX primers – 400 bp (within pCC1BAC-yeast) |
| BK1330_F | GCCATCCCCCTGCTCTACGA | pDEINO1 MPX primers – 650 bp (junction between ori and KatA promoter) |
| BK1330_R | GTGATACTCCGACCAGAGAGACGA | pDEINO1 MPX primers – 650 bp (junction between ori and KatA promoter) |
| pDEINO2 MPX Primers | | |
| BK1327_F | GTGGAGCGGATTATGTCAGCAATG | pDEINO2 MPX primers – 200 bp (pCC1BAC-yeast and 1 kb homology junction) |
| BK1428_R | GCCCCCACCTACGACGTG | pDEINO2 MPX primers – 200 bp (pCC1BAC-yeast and 1 kb homology junction) |
| BK1412_F | GTGGACATTAGTCAGTGGCATCGT | pDEINO2 MPX primers – 300 bp (within DrCm^R^) |
| BK1330_R | GTGATACTCCGACCAGAGAGACGA | pDEINO2 MPX primers – 300 bp (within DrCm^R^) |
| BK1329_F | GCAGTAGCAGAACAGGCCACAC | pDEINO2 MPX primers – 400 bp (within pCC1BAC-yeast) |
| BK1329_R | GGGTAATTCTGCTAGCCTCTGCAA | pDEINO2 MPX primers – 400 bp (within pCC1BAC-yeast) |
| MP1 Megaplasmid MPX Primers | | |
| BK1520_F | GGCAGCTTCAGGGACGTGTC | MP1 MPX Primers– 200 bp |
| BK1520_R | CTCCATGTCTTTCCGCTTGGAGG | MP1 MPX Primers – 200 bp |
| BK1521_F | GTATGGGCCCTGACGGCC | MP1 MPX Primers – 300 bp |
| BK1521_R | GCTGGCCGAACTGGAAGAGG | MP1 MPX Primers – 300 bp |
| BK1522_F | GATCCACCAGGCGAGGGC | MP1 MPX Primers – 400 bp |
| BK1522_R | CTATATCGAGGGCAGCGGCC | MP1 MPX Primers – 400 bp |
| BK1523_F | GATGGACCTGGGAAGCGCC | MP1 MPX Primers – 600 bp |
| BK1523_R | CGGGTTGTGCGTCAATTCGC | MP1 MPX Primers – 600 bp |
| pDEINO29 MPX Primers | | |
| BK2000_F | CGAAACGATCCTCATCCTGT | Nm^R^ |
| BK2000_R | AGGAAGCGGAACACGTAGAA | Nm^R^ |
| BK2003_F | GGCCCACTTCATCACAGAGT | ORF14075 |
| BK2003_R | CCGAACAGGTCCTGGAAGTA | ORF14075 |
| BK2005_F | ACGACCATCACACCACTGAA | pCC1BAC-yeast backbone |
| BK2005_R | CATGACCAGCGTTTATGCAC | pCC1BAC-yeast backbone |
| pDEINO30 MPX Primers | | |
| BK2001_F | CTGATTGTCATCAGCGCATT | Tet^R^ |
| BK2001_R | CAAAGTTGCAGCCGAATACA | Tet^R^ |
| BK2004_F | GCTGGTAAATGCCCTTCGTA | ORF15360 |
| BK2004_R | TCTACGCCGACTTCCTGTTC | ORF15360 |
| BK2005_F | ACGACCATCACACCACTGAA | pCC1BAC-yeast backbone |
| BK2005_R | CATGACCAGCGTTTATGCAC | pCC1BAC-yeast backbone |
| pDEINO32 MPX Primers | | |
| BK2001_F | CTGATTGTCATCAGCGCATT | Tet^R^ |
| BK2001_R | CAAAGTTGCAGCCGAATACA | Tet^R^ |
| BK2005_F | ACGACCATCACACCACTGAA | pCC1BAC-yeast backbone |
| BK2005_R | CATGACCAGCGTTTATGCAC | pCC1BAC-yeast backbone |
| BK2216_F | CCTGACCGAAAGAGAGTTCG | Mrr |
| BK2216_R | GTAGCGGGAGGTCGTCATAA | Mrr |
| pDEINO33 MPX Primers | | |
| BK2000_F | CGAAACGATCCTCATCCTGT | Nm^R^ |
| BK2000_R | AGGAAGCGGAACACGTAGAA | Nm^R^ |
| BK2002_F | GAAGAACTGCCTGAGCGGTA | ORF2230 |
| BK2002_R | GTCCATGCTGCTCTGAAACA | ORF2230 |
| BK2005_F | ACGACCATCACACCACTGAA | pCC1BAC-yeast backbone |
| BK2005_R | CATGACCAGCGTTTATGCAC | pCC1BAC-yeast backbone |
